# Antibody-mediated inhibition of ADAMTS-7 reduces experimental atherosclerosis

**DOI:** 10.64898/2026.08.11.744317

**Authors:** M. Amin Sharifi, Johannes Riechel, Michael Winkler, Tan An Dang, Christian Graesser, Philipp Müller, Carla Abrahamian, Nikita Panyam, Priscilla S. Briquez, Holger Spiegel, Hendrik B. Sager, Nicole Raven, Heribert Schunkert, Thorsten Kessler

## Abstract

**Objective:** One of the strongest genetic associations with coronary artery disease (CAD) risk maps to the metalloproteinase ‘a disintegrin and metalloproteinase with thrombospondin motifs 7’ (ADAMTS-7) locus. The protein was shown to promote plaque formation and instability. We aimed to generate and evaluate an antibody-based strategy targeting ADAMTS-7 therapeutically to reduce atherosclerotic plaque formation.

**Approach and Results:** A truncated form of human ADAMTS-7 was produced in *Nicotiana benthamiana* and used as antigen for antibody generation by hybridoma technology. Eight monoclonal antibodies (mAbs) were screened, among which ADAMTS-7-mAb32 (mAb32) demonstrated the highest affinity, as confirmed by surface plasmon resonance analyses and immunoblotting against full-length ADAMTS-7. *In vitro*, mAb32 inhibited interactions of ADAMTS-7 with its substrates TIMP-1 and SVEP1 in a dose- and time-dependent manner, as assessed by time-resolved Förster resonance energy transfer assays. To assess therapeutic efficacy *in vivo*, *Apoe*^-/-^ mice were fed a Western diet for ten weeks and treated with weekly injections of mAb32 or control IgG over the last six weeks. *En face* aortic Oil Red O staining revealed significantly reduced plaque area in the treatment group, without changes in plasma cholesterol levels or body weight. No evidence of liver or kidney toxicity was observed.

**Conclusion:** Monoclonal antibody-based inhibition of ADAMTS-7 reduced atherosclerotic burden *in vivo* without affecting lipid metabolism, supporting ADAMTS-7 as a viable therapeutic target in CAD. Further development of mAb32 may provide a cholesterol-independent treatment strategy for atherosclerosis.

## Introduction

Ischemic heart disease (IHD) remains the leading cause of death worldwide^1^. While conventional risk factors such as smoking, diabetes, hypercholesterolemia, and hypertension explain a substantial proportion of cardiovascular mortality^2^, a significant component of risk is genetically determined. Family-based and twin studies have long supported a strong heritable component of coronary artery disease (CAD)^3–5^ risk. Genome wide association studies (GWAS) led to the identification of hundreds of genomic variants influencing CAD risk^6,7^. Among the GWAS-identified loci with the strongest effect sizes is *ADAMTS7*^8–10^, which encodes the secreted metalloproteinase ‘a disintegrin with thrombospondin motifs 7’ (ADAMTS-7)^11^ involved in extracellular matrix (ECM) remodeling. ADAMTS-7 was shown to promote vascular smooth muscle cell migration, modulate endothelial cell phenotypes, and influence matrix turnover through substrates such as COMP, TSP1, and TIMP1^12–14^. Polymorphisms at the *ADAMTS7* locus were further associated with high-risk plaque phenotypes^15–17^, while genetic deletion of *Adamts7* in hyperlipidemic mice was shown to reduce atherosclerotic plaque burden and neointima formation following vascular injury^13,18^.

Importantly, loss of ADAMTS-7 function has not been associated with deleterious phenotypes in animal models, rendering it an attractive therapeutic target^13^. Recent attempts to modulate ADAMTS-7 activity, including vaccination strategies, demonstrated reduced atherosclerosis in mice^19^. However, reversible pharmacological tools – such as monoclonal antibodies or small-molecule inhibitors – are currently lacking, or have not made it into clinical application. This may, in part, be due to the absence of suitable screening assays for ADAMTS- 7 activity and an incomplete understanding of which protein domains are most amenable to therapeutic targeting. ADAMTS-7 comprises a catalytic metalloproteinase domain responsible for proteolytic activity and protein interactions, including binding to TIMP-1^14^, as well as multiple C-terminal TSP-1 repeats that contribute to substrate recognition and binding^12,13^, which are interspersed with cysteine-rich and spacer regions^20^. These structural features provide potential sites for antibody-mediated inhibition of ADAMTS-7 function.

In this study, we aimed to develop and characterize monoclonal antibodies targeting ADAMTS-7 to explore their potential as tools within therapeutic strategies to prevent or reduce atherosclerotic plaque formation.

## Methods

*A detailed, expanded Methods section is available in the **Supplemental Material***.

### Production and purification of ADAMTS-7_trunc_

A truncated form of human ADAMTS-7 (ADAMTS-7_trunc_) was generated as an antigen for monoclonal antibody development. The construct comprised the C-terminal region containing thrombospondin type-1 (TSP-1) domains 5–8 (∼43 kDa) followed by a C-terminal His-tag. The corresponding gene sequence was cloned into the binary pTRAkc plant expression vector, a derivative of pPAM (GenBank reference: AY027531), and introduced into *Agrobacterium tumefaciens* for transient expression in *Nicotiana benthamiana* leaves. Leaves were harvested and homogenized in extraction buffer, followed by clarification via centrifugation. Recombinant human truncated ADAMTS-7 (ADAMTS-7_trunc_) protein was then purified using Ni-NTA affinity chromatography and subsequently used as an antigen for monoclonal antibody generation.

Monoclonal antibodies were generated by BioGenes GmbH (Berlin, Germany) using the hybridoma technique as originally described by Köhler and Milstein and further detailed by Holzlöhner and colleagues^21,22^. Briefly, BALB/c mice were immunized intraperitoneally with recombinant human truncated ADAMTS-7 (ADAMTS-7_trunc_). The protein was emulsified in Freund’s complete adjuvant, followed by booster injections with incomplete Freund’s adjuvant at two-week intervals. Mice received a priming dose of 100 µg, followed by two booster injections of 50 µg each and a final pre-fusion boost of 50 µg. Three days after the final boost, spleens were harvested and splenocytes were isolated and fused with SP2/0-Ag14 myeloma cells using polyethylene glycol (PEG 1500; Roche, Basel, Switzerland). Fused cells were seeded into 96-well plates containing hypoxanthine-aminopterin-thymidine selection medium and incubated at 37 °C. Hybridoma supernatants were screened for ADAMTS-7- specific antibody production by enzyme-linked immunosorbent assay (ELISA). Positive clones were expanded and subjected to limiting dilution subcloning to obtain stable monoclonal cell lines. Selected hybridomas were maintained in hypoxanthine-thymidine medium and used for antibody production and characterization.

### Mouse models

Animal experiments were carried out in compliance with the German animal protection laws and conformed to the guidelines of Directive 2010/63/EU and were approved by the local animal care committee (ROB-55.2-2532.Vet_02-18-177). Atherosclerosis-prone mice B6.129P *Apoe^tm1Unc^* (subsequently referred as *Apoe^-/-^*) were purchased from the Jackson Laboratories (Bar Harbor, ME, USA). The mice were provided with unrestricted access to food and water and were kept in a controlled environment with a 12-hour light-dark cycle, a temperature between 20-22 °C, and a humidity level of 45-60%. To induce atherosclerotic plaque formation, *Apoe*^⁻/⁻^ mice were fed a Western diet (21.2% fat, 0.2% cholesterol by weight; Envigo, Indianapolis, IN, USA) for 10 weeks. To evaluate the therapeutic efficacy of the antibody *in vivo*, *Apoe^-/-^* mice were fed a Western diet for 4 weeks to induce early atherosclerotic lesion formation. Animals were then treated with weekly intraperitoneal injections of either the antibody (10 mg/kg) or an isotype-matched control antibody for an additional 6 weeks, while continuing the Western diet. Each treatment and control group consisted of 12 mice, with equal representation of both sexes (6 males and 6 females per group). Mice were randomly allocated to treatment and control groups, and all data from animal experiments were analyzed in a blinded manner. Health was monitored daily using a standardized scoring system, with predefined humane endpoints for mice showing signs of distress, severe barbering, aggression, or weight loss. Mice were harvested under deep general anesthesia, induced in an induction chamber with 4–5% isoflurane in oxygen until complete loss of pedal reflex.

### Statistical analysis

Data are presented as mean ± standard error of the mean (s.e.m.), median and interquartile range (IQR), or violin plots, as appropriate. Outliers were identified and removed using the ROUT method. Distribution of data was analyzed using D’Agostino & Pearson omnibus test or Shapiro-Wilk test, as appropriate. Normally distributed data were analyzed using unpaired t-test for two comparisons or one-way ANOVA for more than two comparisons with appropriate post-hoc tests for multiple comparisons (depicted in the figure legends). Non-normally distributed data were analyzed using Mann-Whitney test for two comparisons or Kruskal-Wallis test for more than two comparisons with appropriate post-hoc tests for multiple comparisons (depicted in the figure legends). All tests were two-sided. Results of distribution analyses, the applied statistical tests, and all experimental details are provided in **Supplemental Table S4**. P-values/adjusted p-values <0.05 were considered statistically significant. GraphPad Prism for macOS, version 10.4.2 (GraphPad, Boston, MA, USA), was used for most analyses unless otherwise stated.

## Results

### Production of ADAMTS7_trunc_ for antibody development

To produce antigen for monoclonal antibody generation, ADAMTS-7_trunc_ was expressed in *Nicotiana benthamiana* via *Agrobacterium tumefaciens*-mediated transformation. The ADAMTS-7_trunc_ protein was purified by Ni-NTA affinity chromatography. Protein purity and expression were assessed by Coomassie staining and immunoblotting using an anti-His antibody. Coomassie staining showed a prominent band at ∼43 kDa, consistent with the expected molecular weight of ADAMTS-7_trunc_, and immunoblotting confirmed the presence of the His-tagged protein at the corresponding size (**Fig. 1B, C**).

**Figure 1:**
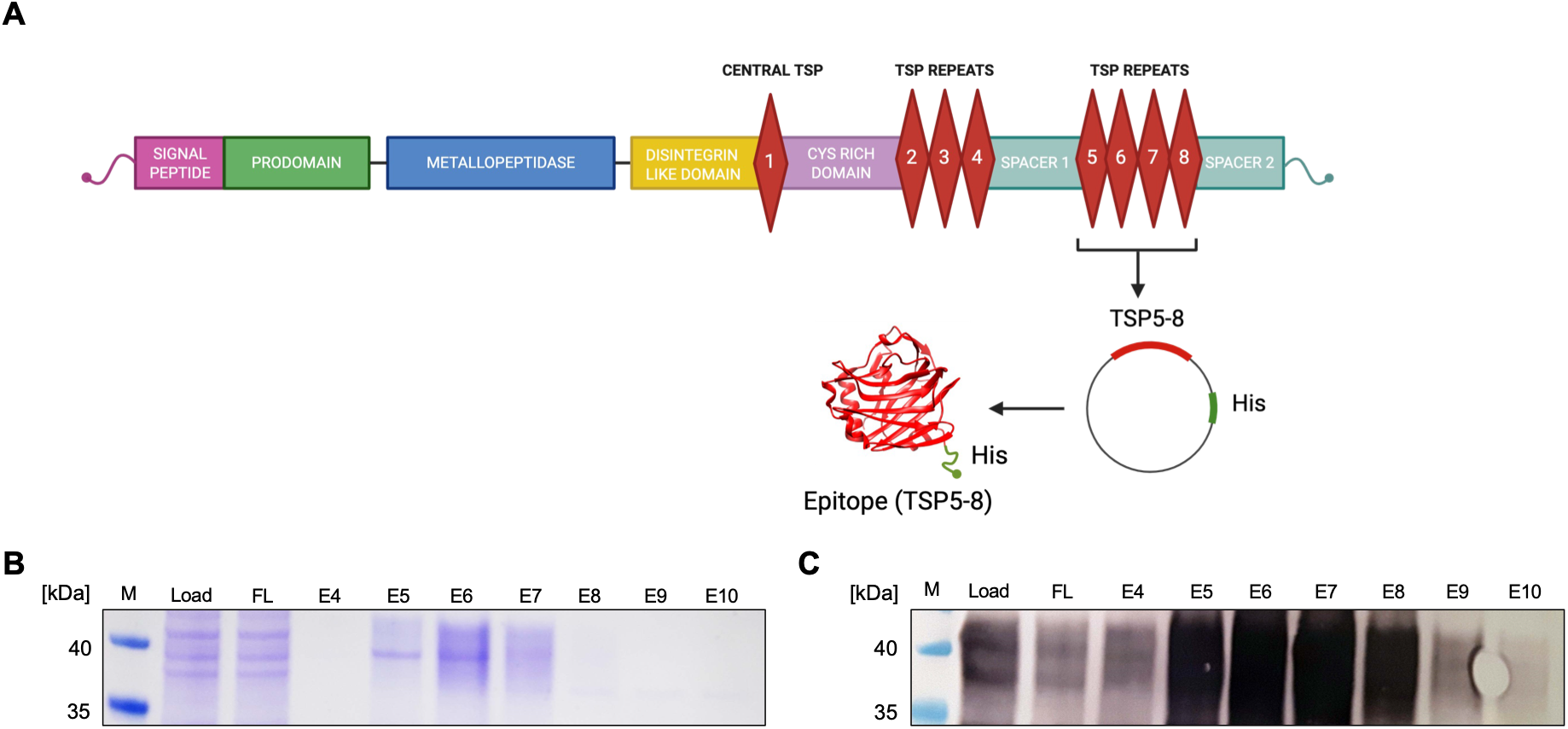
Purification and detection of truncated ADAMTS-7 (ADAMTS-7_trunc_). **A.** We selected a region of ADAMTS-7 containing the fifth to eight thrombospondin repeats (TSP5-8). The sequence encoding the truncated ADAMTS-7 protein (ADAMTS-7_trunc_) was cloned into an expression vector containing a C-terminal His-tag and expressed in *Nicotiana benthamiana* leaves. **B, C.** ADAMTS-7_trunc_- His was purified using Ni-IMAC magnetic beads from leaf material and subjected to Coomassie staining (**B**) and anti-His immunoblotting (**C**). *M*, marker; *FL*, flow-through; *E4–E10*, elution fractions.

### Development and characterization of monoclonal antibodies against ADAMTS-7

To generate monoclonal antibodies against ADAMTS-7 (TS7-mAb), mice were immunized with recombinant ADAMTS-7_trunc_. Splenic B cells from immunized animals were fused with myeloma cells using standard hybridoma technology. Following screening, selection and limiting dilution, eight stable monoclonal hybridoma lines were established for further characterization.

An initial ranking of the eight monoclonal hybridoma lines for ADAMTS-7_trunc_ binding was performed. For this purpose, a mouse-IgG specific capture surface was used to investigate antibody-antigen binding by SPR applying through sequential injections of the antibody and antigen. Comparison of binding rates and endpoint values in relation to the antibody capture signal were determined to identify two candidates (mAb18 and mAb32) with distinctly higher on-rate compared to the other six antibodies.

Because the TSP-repeat region covered by ADAMTS-7_trunc_ shares approximately 85% amino acid sequence homology between human and murine ADAMTS-7 (**Suppl Fig. S1**), we evaluated whether the candidate antibodies could detect full-length ADAMTS-7 of both species, which would enable potential future *in vivo* studies. To this end, human and murine ADAMTS-7 were ectopically expressed in HEK293 cells and analyzed by Western blotting. MAb32 detected clear bands corresponding to the expected molecular weight of full-length ADAMTS-7 in both human and murine samples and showed stronger and more specific detection compared to mAb18 (**Fig. 2B**). While both antibodies were evaluated in subsequent assays, mAb32 consistently demonstrated superior performance. Therefore, the results presented in this study focus primarily on mAb32.

**Figure 2:**
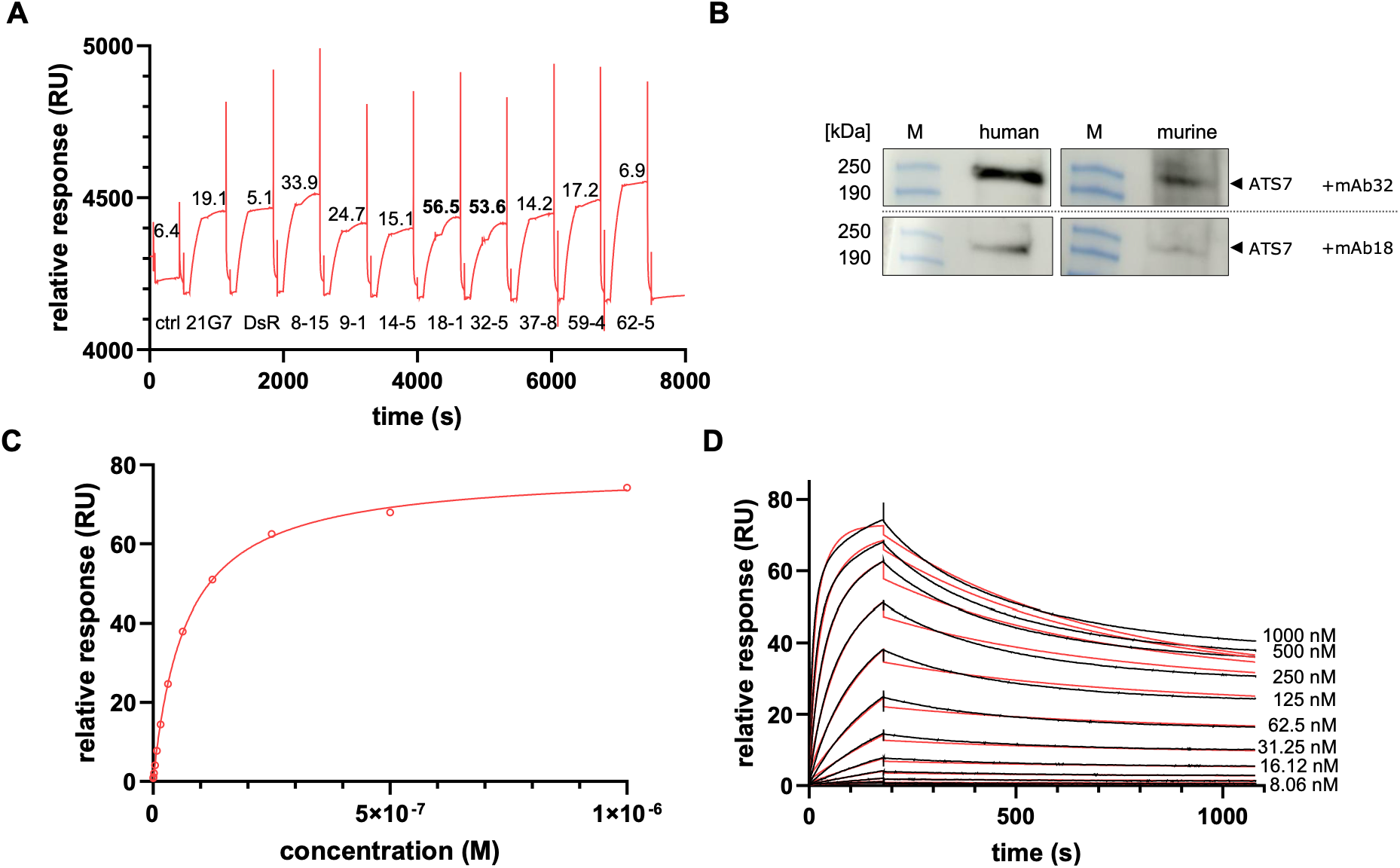
Characterization of monoclonal antibody binding to immobilized truncated ADAMTS- 7. **A.** Surface plasmon resonance (SPR) sensorgram showing the binding responses of the individual generated monoclonal antibodies (mAbs) to immobilized ADAMTS-7_trunc_. Response units (RU) on the y-axis are proportional to the mass of mAb (analyte) bound to ADAMTS-7_trunc_ (ligand). The x-axis represents the progression of time. Each curve represents a distinct mAb interaction, with corresponding RU values indicated above the peaks. Among the tested clones, mAb18 and mAb32 exhibited the highest binding responses to ADAMTS-7_trunc_. **B.** Cell lysates derived from HEK 293 cells expressing full-length human (left) or murine ADAMTS-7 (right) were incubated with mAb32 and subjected to immunoblotting, indicating detection of both human and murine ADAMTS-7. **C.** Steady-state affinity curve for mAb32 binding to ADAMTS-7trunc. Equilibrium responses (RU) are plotted against antibody concentration (M), and data were fitted to a steady-state binding model, yielding an equilibrium dissociation constant (Kd) of 6.766 × 10⁻⁸ M. **D.** SPR sensorgrams depicting binding kinetics of mAb32 to ADAMTS-7trunc at multiple concentrations, fitted with a heterogeneous ligand binding model. RU values over time and the differences in curve shape (red and black) indicate the presence of more than one binding site. *M*, marker.

Binding kinetics of mAb32 were characterized by injecting increasing concentrations of the antibody over immobilized ADAMTS-7_trunc_ using surface plasmon resonance analyses. Sensorgrams revealed distinct association and dissociation phases, with a rapid rise in response units during association and a slow, incomplete return to baseline during dissociation, indicating high-affinity binding with slow off-rate (**Fig. 2C**).

To quantify the interaction, a heterogeneous ligand model was applied. The resulting fit suggested at least two distinct binding interactions, with calculated equilibrium dissociation constants of Kd_1_=7.3·10^-10^ M (high-affinity site) and Kd_2_=7.8·10^-8^ M (lower-affinity site) (**Fig. 2D**). These findings indicate complex binding behavior of mAb32, potentially reflecting interaction with conformationally or spatially distinct epitopes on the truncated antigen surface.

### mAb32 disrupts ADAMTS-7 binding to targets in atherosclerosis

We previously reported that ADAMTS-7 interacts with TIMP-1, potentially contributing to plaque destabilization^14^. To test whether mAb32 can inhibit this interaction, we used a time-resolved Förster resonance energy transfer (TR-FRET)-based assay. Addition of increasing concentrations of mAb32 (0 to 0.05 μg/μl) to the ADAMTS-7/TIMP-1 complex after seven hours of incubation, resulted in a concentration-dependent reduction of the FRET signal (p_trend_=1.22·10^-2^; **Fig. 3A**), with the strongest inhibition observed at 0.05 μg/μl.

**Figure 3:**
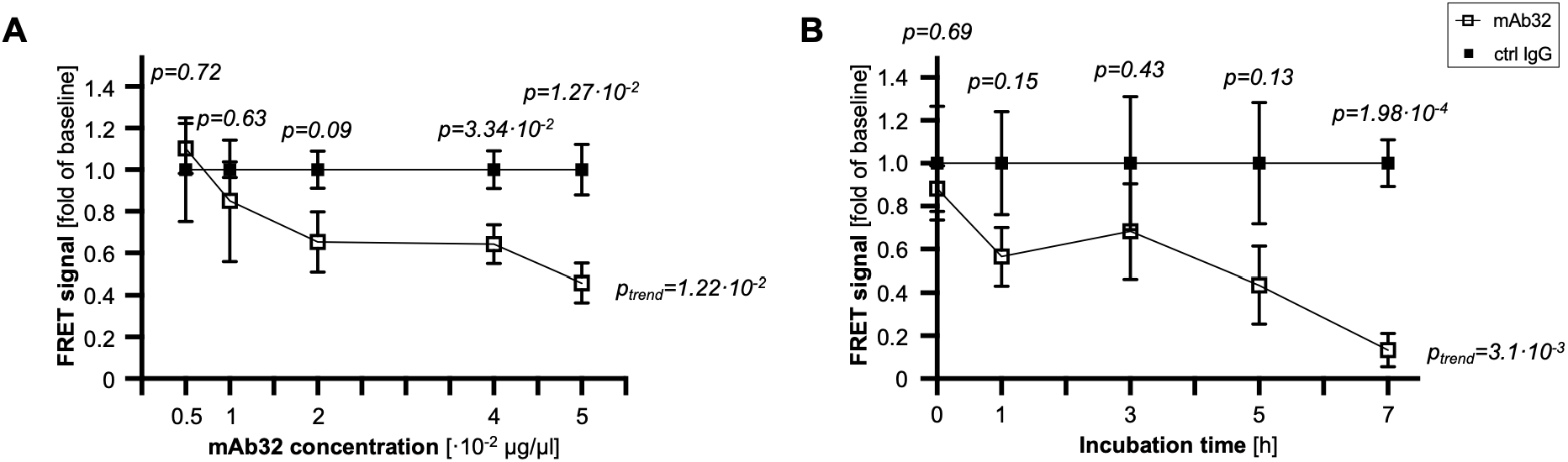
Inhibition of ADAMTS-7’s interaction with TIMP-1 by mAb32. **A.** Concentration- dependent inhibition of the ADAMTS-7–TIMP-1 interaction by mAb32 as indicated by a decrease in the FRET (Δ665/620) signal (n=4 independent experiments). **B.** Time-dependent inhibition of the ADAMTS-7-TIMP-1 interaction by mAb32 (0.05 μg/μl) as indicated by a decrease in the FRET (Δ665/620) signal (n=5 independent experiments). Data are mean and s.e.m. Ordinary one-way ANOVA with post-hoc test for linear trend and Student’s t-test for comparisons of different concentrations/time points.

Using this concentration, we next assessed the time-dependence of inhibition. Over a seven-hour period, mAb32 progressively reduced the FRET signal (p_trend_=3.1·10^-3^; **Fig. 3B**), with maximal effect reached at seven hours, beyond which the signal plateaued.

SVEP1, a recently identified substrate of ADAMTS-7^13^, has itself been genetically linked to CAD risk^23^. To assess whether ADAMTS-7 mediates SVEP1 degradation, HEK293 cells were co-transfected with SVEP1 and either full-length ADAMTS-7, a catalytically inactive variant (ADAMTS-7_Δcat_), or an empty plasmid (mock). Immunoblotting showed reduced SVEP1 levels in the presence of catalytically active ADAMTS-7, indicating substrate cleavage (**Fig. 4A**). Similar findings were observed upon immunoblotting of matrix SVEP1 derived from the supernatant of the same samples, demonstrating a marked reduction in SVEP1 levels in the presence of catalytically active ADAMTS-7 (**Suppl. Fig. S2**).

**Figure 4:**
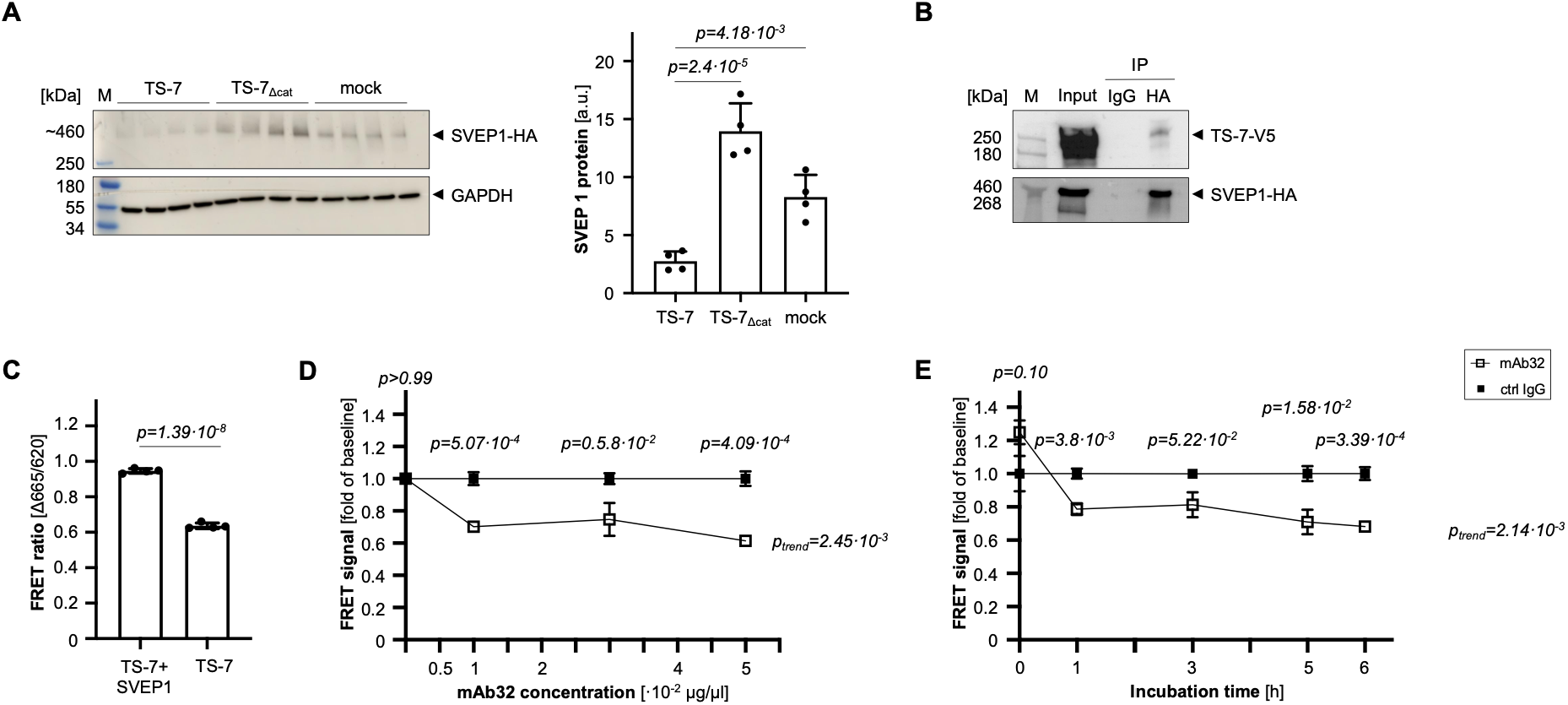
Interaction between ADAMTS-7 and SVEP1. **A.** Degradation assay (left) of SVEP1 in cells co-expressing full-length ADAMTS-7 (TS7), catalytically inactive ADAMTS-7 (TS7_Δcat_), or an empty vector (mock). GAPDH served as a loading control. Quantification (right) revealed higher SVEP1 protein levels in the TS7_Δcat_ and mock groups as compared to full-length ADAMTS-7 (n=4 independent experiments). **B.** Protein-protein interaction between ADAMTS-7 (TS-7-V5) and SVEP1- HA as indicated by co-immunoprecipitation. SVEP1-HA was precipitated using anti-HA antibody-coupled beads, IgG-coupled beads served as control. Immunoblotting of IP fractions confirmed the presence of both ADAMTS-7 and SVEP1 in HA-IP samples, indicating a direct interaction of ADAMTS-7 and SVEP1. **C.** Confirmation of the protein-protein interaction between ADAMTS-7 and SVEP1 using FRET. Quantification of the FRET (Δ665/620) ratio revealed a higher signal when both ADAMTS-7 and SVEP1 were present in comparison to negative control (n=4 independent experiments). **D.** Concentration-dependent inhibition of the ADAMTS-7-SVEP1 interaction by mAb32 as indicated by a decrease in the FRET (Δ665/620) signal (n=4 independent experiments). **E.** Time-dependent inhibition of the ADAMTS-7-SVEP1 interaction by mAb32 (0.05 μg/μl) as indicated by a decrease in the FRET (Δ665/620) signal (n=4 independent experiments). Data are mean and s.e.m. Ordinary one-way ANOVA with Sidak’s multiple comparisons test (A), Student’s t-test (C), and ordinary one-way ANOVA with post-hoc test for linear trend and Student’s t-test for comparisons of different concentrations/time points. *FRET*, Förster resonance energy transfer; *IP*, immunoprecipitation; *M*, marker; *TS-7*, ADAMTS-7.

To investigate whether this degradation results from a direct physical interaction, Co-IP and TR-FRET assays were conducted, confirming a direct interaction between ADAMTS- 7 and SVEP1 (**Fig. 4B-C**). FRET signals were detected only when both proteins were co-expressed.

To test the inhibitory potential of mAb32 on this interaction, increasing concentrations (0 to 0.05 μg/μl) were added to the ADAMTS-7/SVEP1 complex. A concentration-dependent decrease in FRET signal was observed (p_trend_=2.45·10^-3^), confirming alteration of the protein-protein interaction (**Fig. 4D**). Also here, a time-dependent effect was observed (p_trend_=2.14·10^-3^) although the inhibition of the signal was already observed after 1h (**Fig. 4E**).

These data demonstrate that mAb32 inhibits ADAMTS-7-mediated interactions with both TIMP-1 and SVEP1, two substrates with known relevance to plaque formation and CAD pathogenesis. Notably, the magnitude of inhibition differed between the two interactions, potentially reflecting differences in baseline FRET signal strength (**Suppl Fig. S3**).

### In vivo efficacy and safety of mAb32 in a murine model of atherosclerosis

To assess the therapeutic efficacy of mAb32 *in vivo*, *Apoe*^-/-^ mice were fed a Western diet for 4 weeks and subsequently received weekly injections of mAb32 (10 mg/kg) or an isotype-matched control IgG antibody for an additional six weeks (**Fig. 5A**).

**Figure 5:**
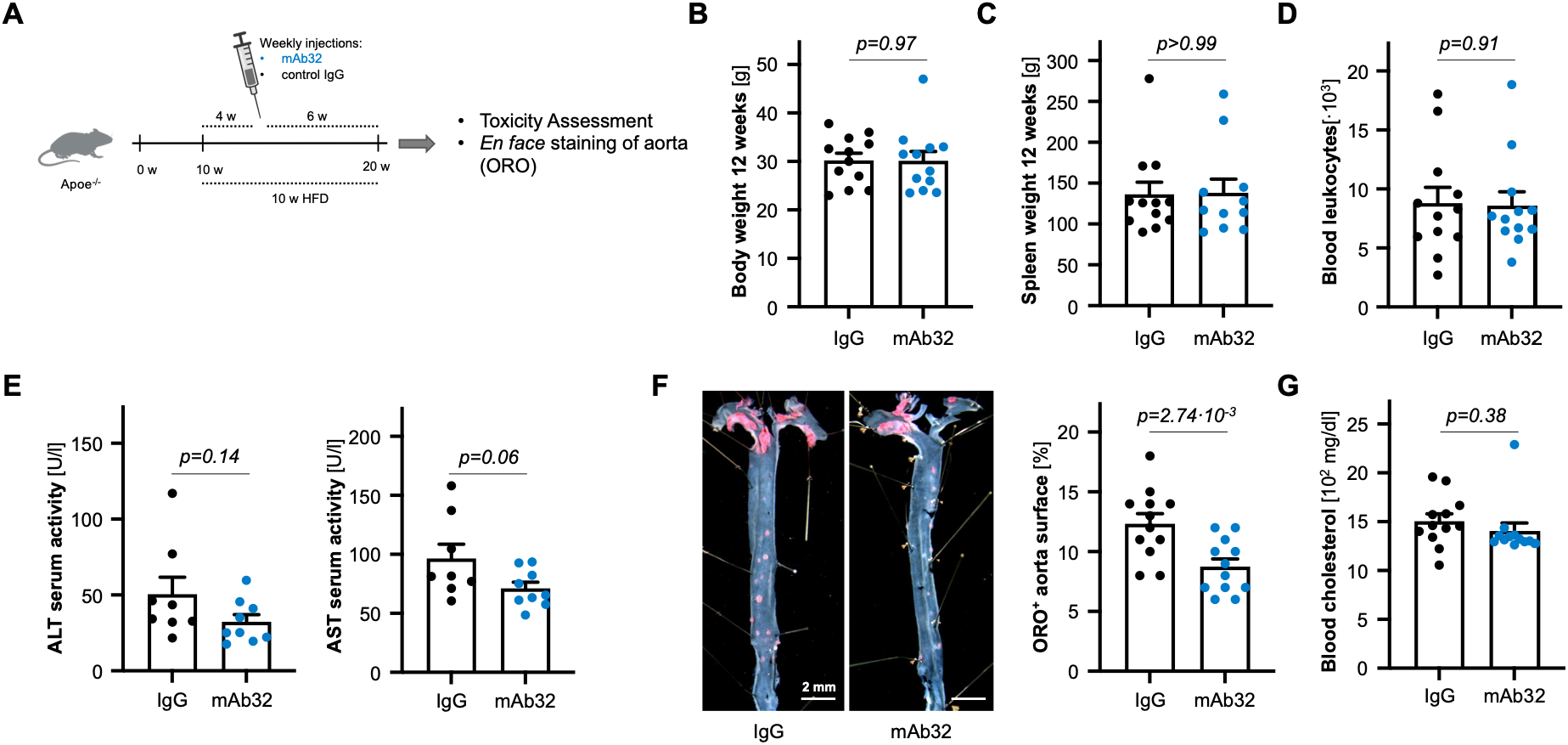
Efficacy and safety of mAb32 under proatherogenic conditions. **A.** Schematic overview of the experimental design. *Apoe^-/-^* mice were fed a Western diet for ten weeks to induce atherosclerotic plaque formation. From week four on, mice received weekly intraperitoneal injections of either mAb32 (10 mg/kg) or an isotype control IgG antibody for six weeks. **B-E.** No differences between treatment groups were observed regarding body weight (**B**) and spleen weight (**C**) at study end, circulating leukocyte count (**D**), or markers of liver function (**E**). **F.** Representative *en face* Oil Red O staining of aortas revealing reduced atherosclerotic plaque burden in mAb32-treated mice as compared with IgG-treated controls (left). *Scale bar: 2 mm*. Quantitative analysis of Oil Red O staining expressed as the percentage of total aortic area covered by plaques indicated a reduction in plaque area compared to control (right). **G.** Total cholesterol levels were comparable between mAb32- and IgG-treated animals. **A-G.** N=12 independent animals per experimental group, and each symbol represents one animal with equal numbers of male and female mice between the groups. Data are mean and s.e.m. Student’s t-test. *ALT*, alanine aminotransferase; *AST*, aspartate aminotransferase; *ORO*, Oil Red O.

Treatment was well tolerated in all animals, with no mortality or signs of systemic toxicity. Body weight (**Fig. 5B**), spleen weight (**Fig. 5C**), total leukocyte counts (**Fig. 5D**), and circulating neutrophils and monocytes (**Suppl. Fig. S4**) did not differ between the groups. Similarly, serum liver enzymes were comparable (**Fig. 5E**).

To assess atherosclerotic plaque burden, we performed *en face* Oil Red O staining of the entire aorta. Consistent with previous reports implicating ADAMTS-7 in aortic plaque development, but rather less in aortic root lesions, quantitative analysis revealed an approximately 30% reduction in plaque area in the mAb32 group compared to control (8.75 ± 0.64 vs. 12.33 ± 0.84% of total aortic surface; p=2.74·10^-3^; **Fig. 5F**). Importantly, total blood cholesterol levels were unchanged between groups (**Fig. 5G**). In mAb32-treated samples, we observed a trend toward increased collagen content and fibrous cap thickness, along with a reduction in necrotic core area, compared to controls. Although these differences did not reach statistical significance, they may suggest a shift toward a more stable plaque phenotype (**Suppl. Fig. S5**). Similar experimental settings were performed with mAb18, with a similar trend but no significant change observed in the overall plaque area (**Suppl. Fig. S6**).

These results demonstrate that systemic inhibition of ADAMTS-7 using mAb32 is safe and reduces atherosclerotic plaque formation *in vivo*, independent of changes in plasma cholesterol levels. This aligns with prior genetic studies suggesting that ADAMTS-7 exerts pro-atherogenic effects through cholesterol-independent mechanisms.

### In silico modeling of ADAMTS-7 interactions with TIMP-1, SVEP1, and mAb32

To complement the experimental findings, we performed *in silico* protein-protein docking using the HADDOCK platform. Docking of ADAMTS-7 with TIMP-1 revealed a multi-domain interaction interface, primarily involving ADAMTS-7’s catalytic and disintegrin-like domains (residues 256–354, 477–480), and the C-terminal region of TIMP-1 (residues 89–181) (**Fig. 6A**). Similarly, docking of the TSP5–8 domain of ADAMTS-7 with the vWF domain of SVEP1 indicated specific contacts between TSP5–6 (Ala1455, Gly1430) and Pro135 on SVEP1 (**Fig. 6B**). Finally, docking simulations using a predicted 3D model of mAb32 revealed that the antibody primarily engages TSP7–8 through its complementarity-determining regions, targeting residues such as Ser1528, Arg1530, and Glu1531 of ADAMTS- 7 (**Fig. 6C**). This aligns with the truncated ADAMTS-7 protein, which was used as the antigen for antibody generation.

**Figure 6:**
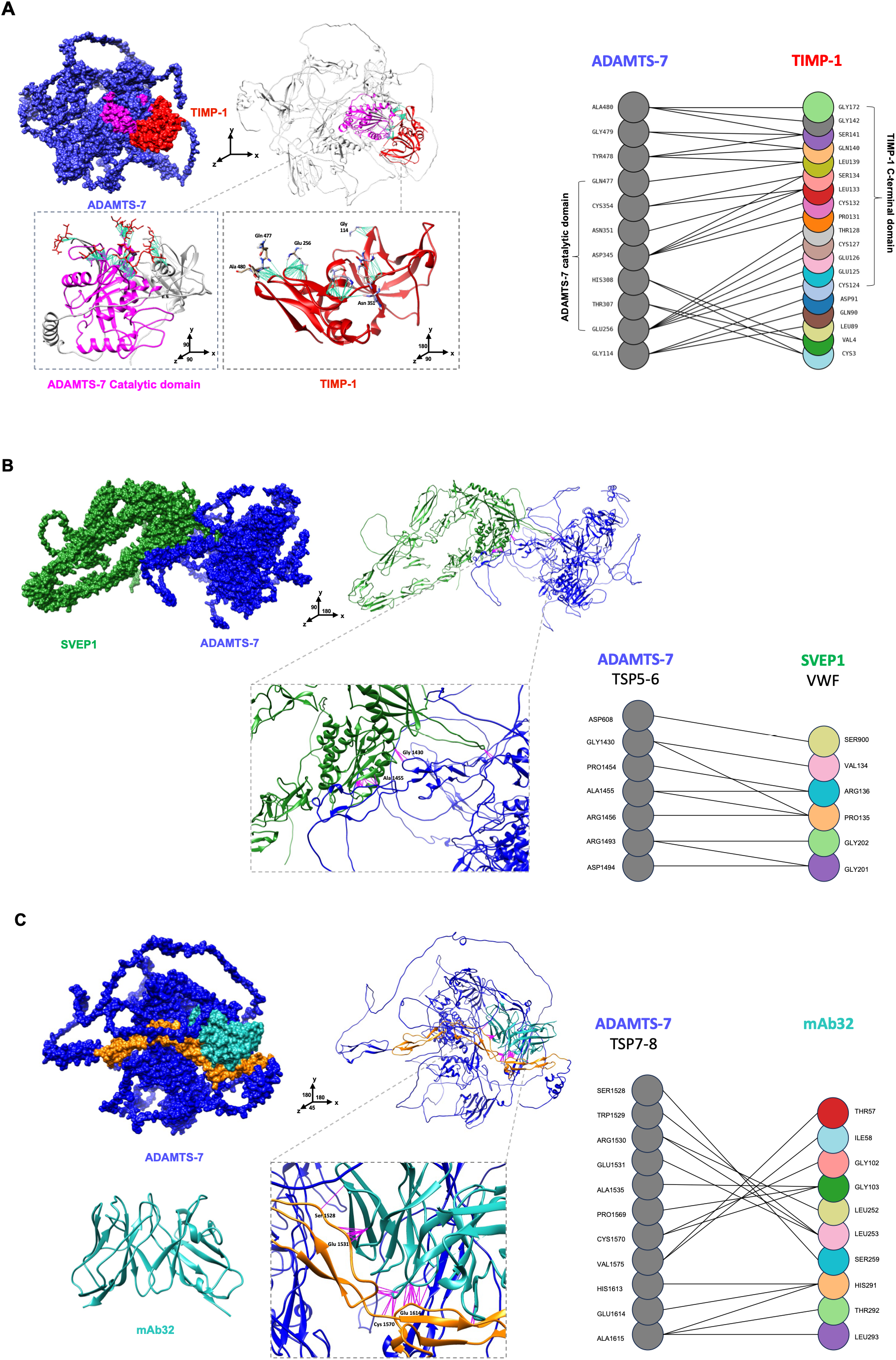
***In silico* docking model and residue-level interaction map of ADAMTS-7 with interaction partners. A.** ADAMTS-7 and TIMP-1 complex. Surface representation shows ADAMTS- 7 (blue) with its catalytic domain highlighted in magenta and TIMP-1 in red at the binding interface. Ribbon and zoomed-in views illustrate the catalytic domain orientation changes and residues of ADAMTS-7 that engage TIMP-1. The right panel displays a residue–residue interaction map, with ADAMTS-7 catalytic domain residues (left, gray) connected to TIMP-1 C-terminal domain residues (right, color-coded) with the key interacting residues. **B.** ADAMTS-7 and SVEP1 complex. ADAMTS- 7 (blue, TSP5–6 domains) engages with SVEP1 (green) at the binding interface, shown in surface and ribbon views with a zoomed inset of interacting residues. The interaction map (right) depicts interaction residues. **C.** ADAMTS-7 and mAb32 complex. The 3D structure of mAb32 was generated using the ABodyBuilder (left). Surface and ribbon representations of ADAMTS-7 (blue) highlight the TSP6–8 domains (orange) and the mAb32 binding region (beige) at the interaction interface. The right panel shows an alternative ribbon view illustrating spatial orientation and epitope engagement. Interactions within 2–4 Å are highlighted by magenta lines, indicating relevant contact sites.

These computational predictions support the specificity of the observed interactions and provide a structural basis for further optimization but require experimental validation.

## Discussion

ADAMTS-7 is a genetic risk factor for CAD and has emerged as a molecular driver of progressive plaque formation and destabilisation^8–10,13,14,18^. By promoting ECM remodeling, ADAMTS-7 impairs vascular plaque stability, making it a compelling target for therapeutic intervention. In this study, we provide a comprehensive evaluation of a monoclonal antibody, mAb32, developed to selectively inhibit ADAMTS-7. We show that mAb32 effectively reduces atherosclerotic plaque burden, likely by disrupting critical protein-protein interactions that drive ECM degradation. mAb32 was generated against a C-terminal ADAMTS-7 epitope, and its binding characteristics were confirmed using SPR, immunoblotting, and immunohistochemistry. SPR revealed high-affinity binding in the nanomolar range, and kinetic profiling indicated a structurally heterogeneous interaction pattern. The antibody cross-reacted with both human and murine ADAMTS-7, confirming their substantial sequence similarity^24^ and supporting its translational potential.

Building on our previously established FRET-based assay^14^, we confirmed that mAb32 disrupts the interaction between ADAMTS-7 and TIMP-1 in a concentration- and time-dependent manner. As TIMP-1 degradation has been associated with increased MMP-9 activity and collagen breakdown, this inhibition might preserve plaque matrix integrity and may promote lesion stability.

We further confirm that SVEP1, an ECM protein genetically and functionally linked to CAD^23,25^, is a direct substrate of ADAMTS-7. *In vitro*, full-length ADAMTS-7 degraded SVEP1, while a catalytically inactive ADAMTS-7 construct did not. Co-IP and FRET assays confirmed a direct interaction. Notably, mAb32 treatment attenuated this interaction in a dose-dependent manner, suggesting that SVEP1 degradation is functionally blocked. While the precise role of SVEP1 in human atherosclerosis requires further clarification, we have previously found that *Svep1* deficiency in *Apoe^-/-^* mice aggravates atherosclerosis through enhanced vascular inflammation and leukocyte recruitment^25^. By inhibiting SVEP1 degradation, mAb32 may thus limit inflammatory cell infiltration and downstream plaque progression.

As described above, HTRF assays revealed that mAb32 effectively disrupts ADAMTS- 7 interactions with both TIMP-1 and SVEP1. A closer examination of the interaction dynamics shows that mAb32 reduced the ADAMTS-7–TIMP-1 FRET signal by up to ∼90%, indicating strong competition between the antibody and TIMP-1 for ADAMTS-7 binding. In contrast, the reduction in FRET signal for the ADAMTS-7–SVEP1 interaction was substantially smaller (∼20–30%), suggesting that SVEP1 is less efficiently displaced by mAb32 under the same assay conditions. This difference was observed despite comparable concentrations of ADAMTS-7 and mAb32 in both assays and an excess of TIMP-1 relative to ADAMTS-7 (molar ratio ADAMTS7:mAb32:TIMP1 ≈ 1:1.2:6.4), whereas SVEP1 was present at a lower relative stoichiometry (ADAMTS7:mAb32:SVEP1 ≈ 1:1.2:0.5). Together with the higher baseline FRET signal observed for the ADAMTS-7–SVEP1 complex, these findings are consistent with a stronger interaction between ADAMTS-7 and SVEP1 than with TIMP-1, potentially reflecting differences in binding affinity under the assay conditions tested. Importantly, because these measurements were performed using a competition-based TR-FRET assay rather than a formal equilibrium binding analysis, the data should not be interpreted as precise affinity estimates but instead as evidence of differential competitive displacement.

*In silico* docking analyses suggested spatially distinct binding interfaces on ADAMTS- 7: TIMP-1 localized near the catalytic domain, SVEP1 bound to TSP5–6, and mAb32 primarily engaged TSP7–8. While these sites do not overlap directly, the observed inhibition of TIMP-1 and SVEP1 interactions suggests that mAb32 may exert allosteric effects by restricting conformational dynamics, altering domain flexibility, or modifying local electrostatics – mechanisms well documented in antibody-mediated modulation of enzymatic proteins^26–28^. Structural studies will be required to validate this predicted binding mode.

In a murine model of atherosclerosis, mAb32 administration significantly reduced plaque burden without affecting cholesterol levels, highlighting a cholesterol-independent mechanism of vascular protection. This is consistent with previous reports on ADAMTS-7’s non-lipid-mediated effects^19^. The antibody was well tolerated, with no evidence of hepatotoxicity, leukocyte abnormalities, or weight loss, reinforcing its therapeutic potential. Beyond its efficacy and safety, antibody-based inhibition of ADAMTS-7 may offer distinct advantages over small-molecule approaches. While small-molecule inhibitors are typically easier to administer and synthesize, achieving selectivity within the ADAMTS family remains challenging due to the high degree of structural homology among their catalytic domains^29^. In contrast, monoclonal antibodies can be designed to target conformational epitopes outside the active site, such as substrate-binding or regulatory domains, thereby increasing specificity and minimizing off-target effects^30^. In this context, mAb32’s ability to bind the TSP7–8 region – rather than the catalytic domain – may help avoid interference with related ADAMTS proteases, while still blocking critical substrate interactions. Moreover, the long serum half-life and defined pharmacokinetics of monoclonal antibodies allow for sustained target engagement, which may be particularly advantageous in chronic vascular conditions such as atherosclerosis. These results support mAb32 as a promising candidate for targeted inhibition of ADAMTS-7. Its mode of action – blocking specific protein–protein interactions while avoiding systemic metabolic effects – could make it particularly useful in patients with residual inflammatory risk, despite lipid-lowering therapy. Additionally, it has been shown that *Adamts-7* expression in mice is transient, occurring mainly during the early stages of atherosclerosis development, whereas its human counterpart appears to be expressed during later stages as well^18^. This suggests that our monoclonal antibody could have an even more pronounced therapeutic effect in humans. While monoclonal antibodies offer high target specificity and long serum half-life, further development steps will be required to advance mAb32 toward clinical application. These include humanization, pharmacokinetic optimization, and improved delivery to vascular lesions.

We wish to acknowledge certain limitations of this study. First, although we demonstrate clear *in vitro* and *in vivo* efficacy of mAb32 in murine models, we did not assess pharmacokinetics, biodistribution, or long-term safety in larger animal models or humans. Second, the precise epitope targeted by mAb32 has not been experimentally mapped, and the proposed binding interface is based on *in silico* modeling; further structural analyses (e.g., cryo-EM or epitope binning) are needed for validation. Third, we focused on TIMP-1 and SVEP1 as primary substrates, but additional ADAMTS-7 targets may contribute to its pro-atherogenic effects and should be explored in future work. Fourth, antibody treatment was administered in a preventive setting, and its efficacy in established lesions or regression models remains to be evaluated. A further limitation is the lack of a direct enzymatic activity assay for ADAMTS-7. Since no validated biochemical substrate assay is currently available, we assessed antibody function using complementary protein–protein interaction assays, substrate degradation experiments, and *in vivo* efficacy.

In summary, these findings establish ADAMTS-7 as a druggable therapeutic target and provide proof-of-concept that antibody-mediated inhibition of ADAMTS-7 attenuates atherosclerosis independently of cholesterol lowering.

## Acknowledgements

The Graphical Abstract was generated using Biorender. Generative AI (ChatGPT-5.0, OpenAI) was used for language refinement and proofreading.

